# Nucleocapsid vaccine elicits spike-independent SARS-CoV-2 protective immunity

**DOI:** 10.1101/2021.04.26.441518

**Authors:** William E. Matchett, Vineet Joag, J. Michael Stolley, Frances K. Shepherd, Clare F. Quarnstrom, Clayton K. Mickelson, Sathi Wijeyesinghe, Andrew G. Soerens, Samuel Becker, Joshua M. Thiede, Eyob Weyu, Stephen O’Flanagan, Jennifer A. Walter, Michelle N. Vu, Vineet D. Menachery, Tyler D. Bold, Vaiva Vezys, Marc K. Jenkins, Ryan A. Langlois, David Masopust

## Abstract

Severe acute respiratory syndrome coronavirus 2 (SARS-CoV-2) is responsible for the COVID-19 pandemic. Neutralizing antibodies target the receptor binding domain of the spike (S) protein, a focus of successful vaccine efforts. Concerns have arisen that S-specific vaccine immunity may fail to neutralize emerging variants. We show that vaccination with HAd5 expressing the nucleocapsid (N) protein can establish protective immunity, defined by reduced weight loss and viral load, in both Syrian hamsters and k18-hACE2 mice. Challenge of vaccinated mice was associated with rapid N-specific T cell recall responses in the respiratory mucosa. This study supports the rationale for including additional viral antigens, even if they are not a target of neutralizing antibodies, to broaden epitope coverage and immune effector mechanisms.

## Introduction

Vaccine candidates targeting SARS-CoV-2 S protein, which is essential for cell entry, were designed based on viral sequences reported in January 2020. Studies of the highly efficacious Pfizer/BioNTech and Moderna mRNA vaccines show a strong correlation between neutralizing antibodies (NAb) response and protection from disease (1, 2). As the virus continues to spread globally, variants have emerged that evade neutralization by vaccine elicited S antibodies (3), which may require continued vaccine adaptation and boosting.

T cells contribute to immunity against respiratory pathogens, including serological variants of influenza virus(4). Unlike NAb, T cell immunity is not limited to surface antigens and immunodominant epitopes vary considerably between individuals due to recognition in the context of genetically diverse MHC molecules. Accordingly, T cell immunity may be less vulnerable to immune selection pressure and viral escape mutations. We immunized rodents against SARS-CoV-2-N to address whether protection could be conferred independently of spike-specific responses (and thus NAb).

## Results

### Ad5-N vaccine confers protection against SARS-CoV-2 infection

Outbred Syrian hamsters are permissive to SARS-CoV-2 infection. We vaccinated hamsters intravenously (IV) with a human adenovirus serotype 5 (Ad5) vector expressing the N sequence (Ad5-N) derived from USA-WA1/2021 strain (WA) or a control Ad5 vector lacking a SARS-CoV2 sequence (Ad5-NULL). 7-8 weeks later, animals were challenged intranasally with 6.8×10^4^ pfu WA SARS-CoV-2. Vaccination reduced weight loss (Fig. 1A, B). Vaccinated hamsters were also challenged with WA or a variant strain B.1.1.7, which contains two amino acid substitutions in N that do not occur within the N_219-227_ immunodominant epitope that we previously defined in C57Bl/6 mice (5). Vaccination elicited a significant 30-fold (WA) and 12-fold (B.1.1.7) reduction in median lung viral titer three days following challenge, though the latter was not significant. We next immunized transgenic K18-hACE2 (K18) mice that are highly susceptible to SARS-CoV-2 (6). Ad5-N vaccinated K18 mice experienced significantly reduced weight loss (Fig. 1D, E) and mortality, with 75% surviving challenge vs. 0% in the Ad5-NULL vaccinated group (Fig. 1F) upon challenge with 300 pfu WA SARS-CoV-2.

**Figure 1.**
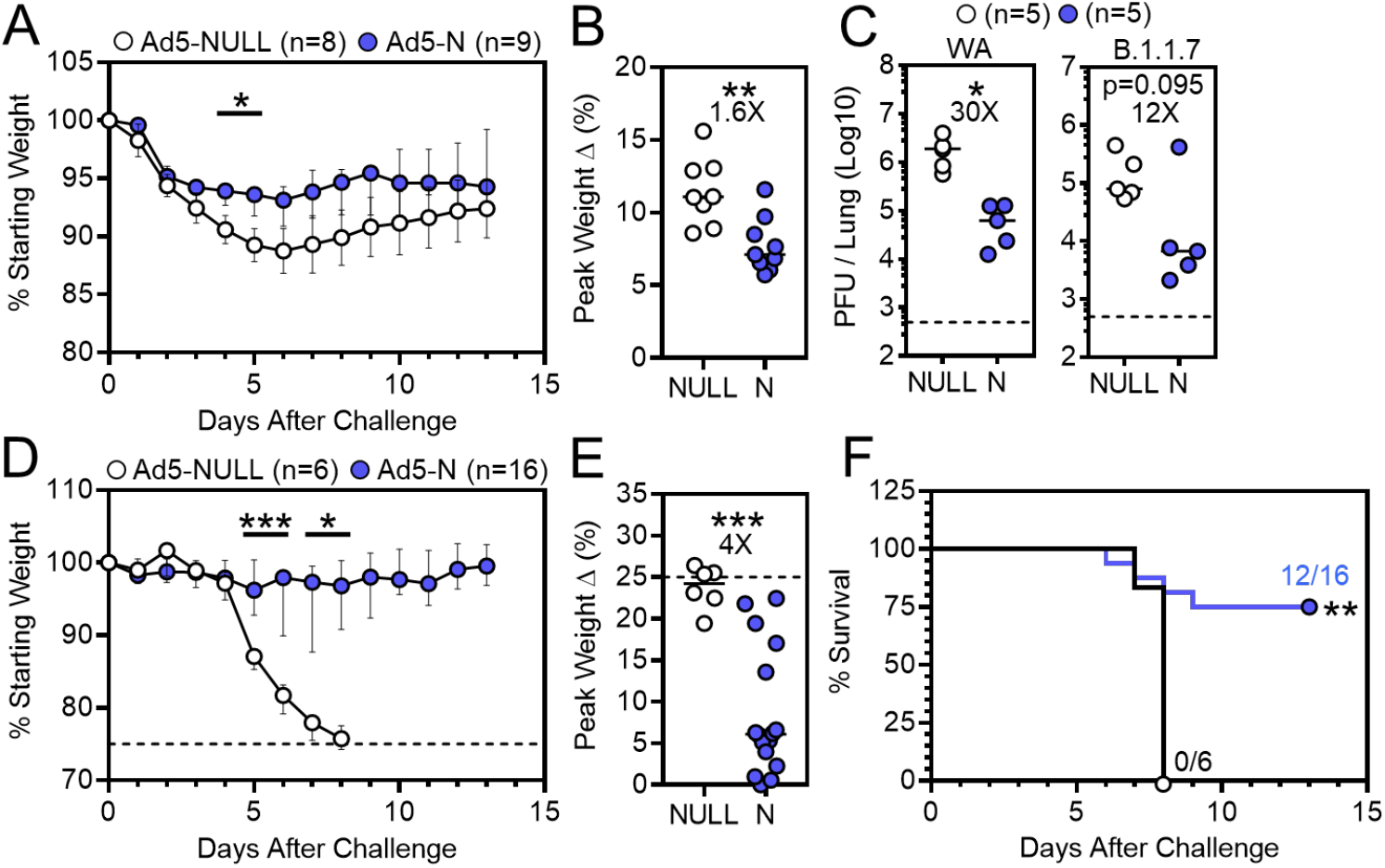
Ad5-N vaccine confers protection against SARS-CoV-2 infection. **A**) Median and **B**) peak weight change after vaccinated hamsters were challenged 7-8 weeks later with WA SARS-CoV-2. **C**) Lung viral titers three days post-challenge with WA (*left*) or B.1.1.7 (*right*) SARS-CoV-2. **D**) Median and **E**) peak weight change (dashed lines represent humane endpoint), and **F**) survival of vaccinated K18-hACE2 mice following intranasal WA SARS-CoV-2 challenge. Bars denote median and interquartile range *, *P* < 0.05; **, *P* < 0.01; ***, *P* < 0.001. Individual comparisons were analyzed using two-sided Mann–Whitney tests (**A-E**) using Bonferroni-Dunn correction for multiple comparisons (**A, D**) and survival by the log-rank test (**F**).

### Memory T cells respond to SARS-CoV-2 challenge

Vaccination established SARS-CoV-2-N_219-227_-specific memory CD8 T cells that persisted in lung, airways, and spleen from 40 to 86 days, and increased in lung draining mediastinal lymph node (Fig. 2A), which may result from accumulation of resident memory T cells (7). We examined whether T cells in vaccinated mice participated in the response four days after WA SARS-CoV-2 challenge. Vaccinated mice showed a ~3.8-log10 reduction in lung viral load (Fig. 2B). Combined antibody-mediated CD4 and CD8b depletion prior to challenge partially abrogated protection, although T cell depletion was not absolute in lungs and elevated N_219-227_-specific CD8 T cell responses were still observed in vaccinated mice (Fig. 2B and data not shown).

**Figure 2.**
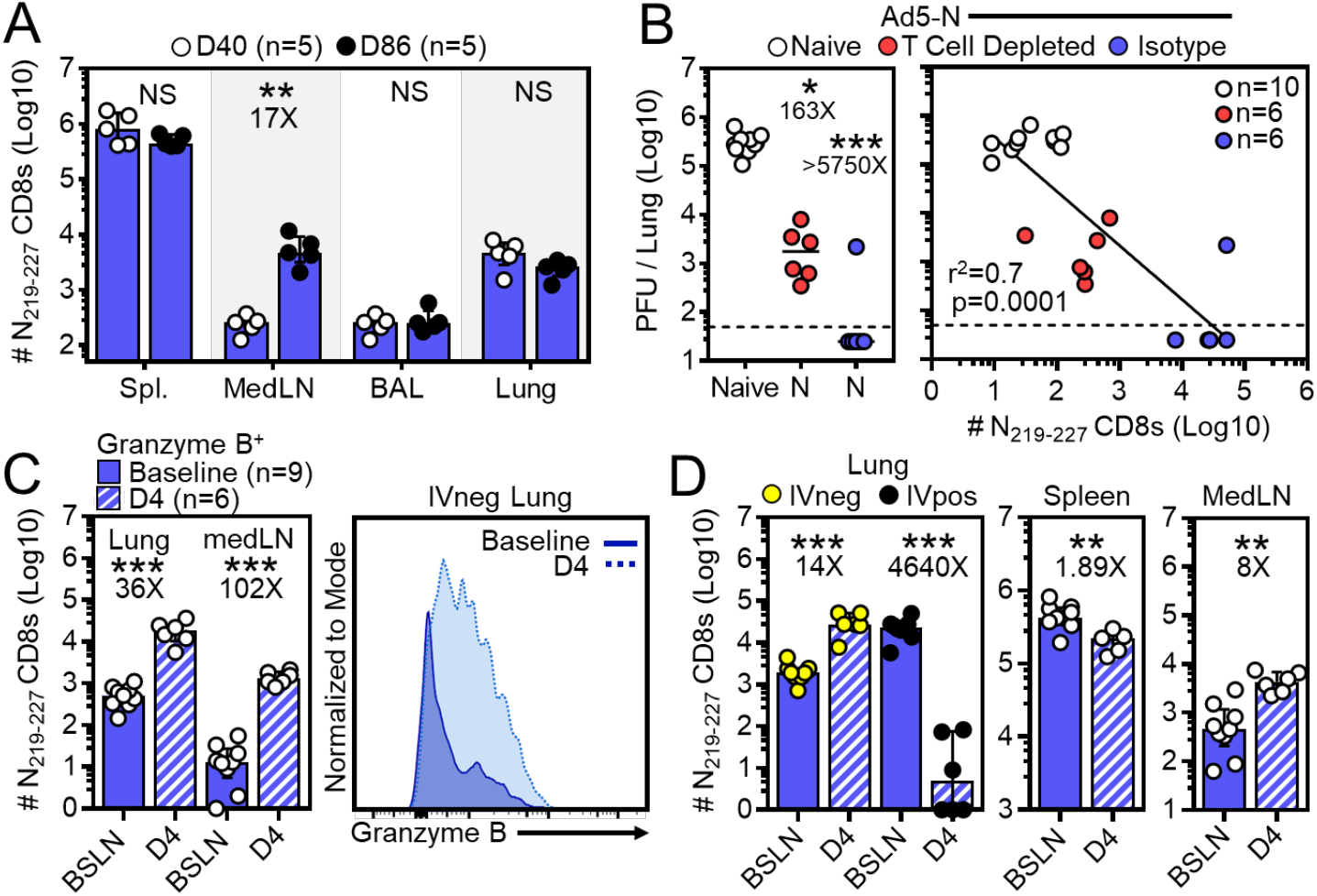
Memory T cells respond to SARS-CoV-2 challenge. **A**) Number of N_219-227_-specific CD8 T cells 40 to 86 days after IV Ad5-N vaccination. **B**) Four days after WA SARS-CoV-2 challenge, viral titers in lungs of naive and Ad5-N vaccinated K18 mice ± T cell depletion (*left*) and viral titers plotted against N_219-227_-specific CD8 T cells within IV^neg^ lung (*right*). **C**) Abundance of granzyme B+ N_219-227_-specific CD8 T cells in IV^neg^ lung or mediastinal LN and **D)** of total N_219-227_-specific CD8 T cells within indicated compartments immediately prior (baseline) and four days after challenge. Bars denote median and interquartile range. *, *P* < 0.05; **, *P* < 0.01; ***, *P* < 0.001 as determined by a two-sided Mann–Whitney test (**A-D**).

Intranasal challenge increased granzyme B+ N_219-227_-specific CD8 T cells in the pulmonary mucosa (Fig. 2C). Total N_219-227_-specific CD8 T cells substantially increased in both lung parenchyma (IVneg, defined by the absence of intravascular anti-CD8a staining, as previously described (8)) and draining mediastinal lymph node, moderately decreased in spleen, and substantially decreased in the lung vasculature (IVpos, Fig. 2D). These data indicate that vaccine-elicited memory CD8 T cells underwent rapid reactivation and migration to the site of viral challenge, and that T cells may have contributed to viral control.

## Discussion

The rapidity and success of SARS-CoV-2 vaccine development and deployment is unprecedented. Most strategies, including vaccines licensed for emergency use in the USA, rely solely on the viral spike. Spike is a logical choice because it contains the receptor binding domain that is the main target of NAb. Nevertheless, SARS-CoV-2 is likely to become endemic. Viral variants have emerged that escape vaccine-elicited NAb, and SARS-CoV-2 may continue to evolve with or without selection pressure from increased global immunity. This study provides evidence in rodents that immunological memory to additional antigens could provide protection, which may involve memory T cells or non-neutralizing antibodies. It appears likely that viral evolution will necessitate vaccine evolution and booster immunizations that address emergent variants. This study supports the rationale for including additional viral antigens into future vaccine candidates to broaden epitope diversity and protection while limiting opportunities for viral escape.

## Materials and Methods

Rodent vaccination, SARS-CoV-2 challenges, assessment of viral titer and T cell responses and other methods are described in extended methods.

## Acknowledgments

We thank the Biosafety Level 3 Program at the University of Minnesota. Supported by the Office of the Dean of the University of Minnesota Medical School and UMN-Mayo Partnership for Biotechnology and Medical Genomics. SARS-CoV-2, Isolate USA-WA1/2020 and the B.1.1.7. variant were obtained through WRCEVA at UTMB. W.E.M., V.J., J.M.S., S.W., and J.M.T. were supported NIH T32 HL007741, the Canadian Institutes of Health Research Fellowship, NIH T90 DE 022732, F30 DK114942 and NIH T32 AI055433 respectively.

## Author Contributions

W.E.M, V.J., J.M.S., S.W., V.V., M.K.J., R.A.L, and D.M. designed research; W.E.M, V.J., J.M.S., F.K.S., C.F.Q, C.K.M, S.W., A.G.S., S.B., E.W., and S.O. performed research; J.M.T., J.A.W.,M.V., V.D.M., and T.D.B. contributed reagents/analytic tools; W.E.M, V.J., J.M.S., F.K.S., R.A.L, and D.M. analyzed data; W.E.M, V.J., J.M.S., F.K.S., R.A.L, and D.M. wrote paper.

