## Extended Methods for "Nucleocapsid vaccine elicits spike-independent SARS-CoV-2 protective immunity"

*Hamster immunizations and SARS-CoV-2 infection*

Male Golden Syrian Hamsters (Charles River Laboratories) were maintained in BSL-2 containment facility under speciﬁc-pathogen-free conditions at the University of Minnesota in accordance with the Institutional Animal Care and Use Committees guidelines.

Eighteen- to nineteen-week-old hamsters were anesthetized with isoflurane and immunized using a single intravenous (IV) dose of 1.8x10^11 viral particles (vp) replication-deficient HAdV-5 expressing SARS-CoV-2-N (Ad5-N) or a control HAdV-5 vector (Ad5-NULL). The construction of Ad5 vectors was described elsewhere (1). Hamsters were moved to BSL-3 containment for SARS-CoV-2 challenge studies. Animals were anesthetized with isoflurane and challenged intranasally (IN) with a total of 6.75x10^5 PFU in a 100ul volume of either the 2019-nCoV/USA_WA1/2020 (WA) strain or the B.1.1.7 variant SARS-CoV-2/human/USA/CA_CDC_5574/2020 (provided by World Reference Center for Emerging Viruses and Arboviruses at the University of Texas Medical Branch). Animals were weighed and checked daily for changes in their condition. All weight loss studies were terminated thirteen days after the challenge. Hamsters challenged for the determination of viral titers in the lungs were euthanized three days after challenge.

*Mouse immunizations and SARS-CoV-2 infection*

Six- to ten-week-old female B6 or hemizygous K18.hACE2 mice (stock no. 034860, Jackson Laboratories) were maintained in speciﬁc-pathogen-free conditions at the University of Minnesota in accordance with the Institutional Animal Care and Use Committees guidelines.

Mice were immunized IV with 5x10^10 viral particles of Ad5-N vector or Ad5-Null.

Immunized K18.hACE2 mice were challenged with SARS-CoV-2 in the BSL-3 containment facility. Animals were anesthetized with ketamine/xylazine, laid supine, and challenged intranasally with 300 PFU of WA SARS-CoV-2 in a 30 ul volume. Infected animals were monitored daily for health, weight loss and morbidity and were deemed terminal if they reached 75% of starting weight or were found dead at the daily inspection. All weight loss experiments were terminated 13 days after infection. Mice assigned for flow cytometry analysis and the concurrent determination of viral titers in the lungs were euthanized four days after challenge.

Intravascular labeling, cell isolation, and ﬂow cytometry

To discriminate extravascular cells from intravascular cells, mice were injected IV with BV605-conjugated anti-CD8a Ab, sacriﬁced after 3 min, and tissues were harvested(1). Mouse lungs were mechanically minced into small pieces and further dissociated in PBS using gentlemacs tubes. For SARS CoV-2 challenge studies, dissociated tissue was split equally into two parts, one for quantification of SARS CoV-2 viral titers and the other for isolation of lymphocytes for flow cytometric analyses. Isolation of cells for flow cytometric analysis was performed as previously described (2). Cells were stained with Abs to CD8a (53-6.7), CD62L (MEL-14), Ly6C (HK1.4), CD44 (IM7), CD69 (H1.2F3), and CD103 (M290) (all from BD Biosciences, Tonbo Biosciences, BioLegend, or Affymetrix eBiosciences) and H2-D^b^/N_219-227_ MHC class I tetramers (made in-house) and ghost dye 780 (Tonbo Biosciences). For monomer preparation, the N_219-227_ peptide sequence was LALLLLDRL. Stained samples were acquired on LSRII or LSR Fortessa ﬂow cytometers (BD Biosciences) and analyzed with FlowJo software (BD Biosciences). Cells were gated on singlets, live lymphocytes, CD8 T cells, and/or tetramer-speciﬁc cells as indicated.

Hamster and mouse lung lysate preparation

Hamsters and mice were euthanized 3 and 4 days after SARS-CoV-2 infection, respectively.  Lungs were harvested from the hamsters, weighed, and homogenized in PBS using gentleMACS M tubes and a gentleMACS dissociator on the protein setting. Mouse lungs were weighed and processed as described above before being further homogenized in PBS using gentleMACS M tubes. Tubes were centrifuged to pellet debris. The supernatants were collected, aliquoted, and stored at -80°C.

Cell Culture and Plaque Assays

Vero E6 cells were maintained in Dulbecco’s modified Eagle medium (DMEM) with 10% fetal bovine serum (FBS) and 1% penicillin-streptomycin (pen-strep). Infections were carried out in an infection medium (DMEM with 1% pen-strep, 1% Non-Essential Amino Acids, 10mM HEPES, and 2% FBS).

The plaque assay was adapted from Mendoza et al(3). Ten-fold dilutions of the homogenized lung samples were adsorbed to Vero E6 cells (ATCC® CRL‐1586™). The cells were incubated for 1 hour at 37°C in a 5% CO_2_ incubator, rocking every 10 minutes. Cells were washed with PBS and overlaid with a carboxymethylcellulose (CMC) overlay media (MEM, 1.5% CMC) and incubated at 37°C in a 5% CO_2_ incubator for 3 days. The overlay media was removed, cells were washed with PBS, and fixed for 1 hour with 4% paraformaldehyde at room temperature. The fixative was discarded, and the cells were stained with crystal violet (0.5%) for 5-15 minutes. Cells were washed with distilled water 2-4 times, blotted dry, and plaques were enumerated.
